# Developing *in vitro* patient derived CERvical Cancer OrganoidS (CERCOS) as a potential preclinical tool in cervical cancer research

**DOI:** 10.1101/2025.06.19.660648

**Authors:** Surbhi Singla, Rashmi Bagga, Radhika Srinivasan, Prateek Bhatia, Shalmoli Bhattacharyya

## Abstract

Advanced cervical cancer remains a major cause of mortality in women worldwide as it has limited treatment options and recurrence is very common. This highlights the necessity to develop patient derived organoids (PDOs) as preclinical models, that can recapitulate the clinical heterogeneity of the cancer in terms of molecular features, and genetic background. The PDOs have potential for guiding personalized treatment in clinical practice. In this study, we have established patient-derived cervical cancer organoids or CERvical Cancer OrganoidS (CERCOS) from biopsy samples of five patients having two different subtypes (squamous cell carcinoma and adenocarcinoma) using a modified protocol. The CERCOS were characterized to assess their genetic and phenotypic similarity with the parent tissue. The organoids developed *in vitro* showed preservation of the histopathological and somatic mutational profile of the parent tissue. Expression of cervical cancer-related genes, including PIK3CA, MET, and LRP1B, was found to be comparable between the organoids and the parent tissue. Moreover, characterization of the PDOs post cryopreservation showed the maintenance of the morphological and histopathological features of the parent tissue. This study demonstrates that the CERvical Cancer OrganoidS (CERCOS), established using the current protocol, preserves genetic and phenotypic similarity and can serve as a platform for precision medicine and biobanking purposes.

## Introduction

Cervical cancer has been a global threat due to its high recurrence rate. Patients often develop resistance to chemo- and radio-therapy which causes tumor relapse and some patients do not respond to these therapies at all. The response of the patients to therapy is quite unpredictable and by the time, it becomes evident, the patient has already progressed to the advanced stage. Many drugs that show promising results during laboratory studies fail during clinical trials. This shortcoming can be attributed to the fact that many cancer models poorly recapitulate the patient’s tumor which hinders the translation of scientific knowledge from bench to bedside. In the last few decades, organoid technology has emerged as a powerful tool with broad application prospects in basic cancer research. Patient-derived organoids (PDOs) are known to replicate the genetic, phenotypic, and functional characteristics of the parent tumor, making a highly relevant model in advancing personalized medicine [1]. As a functional model that can mimic the multi-omics characteristics and drug sensitivity of the original tumor, PDOs can be employed as an in vitro substitute for novel drug development. PDOs have been successfully established and used to predict the chemotherapy or targeted therapy response in various cancers including breast cancer, pancreatic cancer, lung cancer, rectal cancer and ovarian cancer [2–11]. PDOs provide a platform for customization of cancer treatment for precision/personalized medicine by taking individual heterogeneity into account. This can improve the therapeutic efficacy with the least drug side effects and reduce the treatment costs.

In this study, we have developed cervical cancer PDOs as representative preclinical in vitro tool which can recapitulate the molecular and phenotypic landscape of the original cervical tumor. Earlier, Villa et al. had established organoids from isolated human cervical keratinocytes using organotypic raft culture method [12]. This method involved establishment of keratinocyte cell line from the patient tissue followed by the generation of organoids from the cell line. This was relatively a time-consuming method and the cells tend to lose their genetic heterogeneity during 2D culture. Later, Maru et al. (2019) established organoids from a single patient of clear cell carcinoma using modified Matrigel Bilayer Organoid Culture (MBOC) protocol [13]. In this method, the organoids were sandwiched between two layers of matrigel which was reported to be highly efficient method of organoid formation for gynecological tumors. Recently, Lohmussaar et al. (2021) and Seoul et al (2022) reported organoid formation by using Matrigel dome method in which organoids were trapped inside Matrigel drops [14–15]. All these groups have utilized different protocols and different media composition for cervical cancer organoid establishment. In the present study, cervical cancer organoids have been established using a modified protocol and have been extensively characterized. Moreover, it is important to cryopreserve the organoids without loss of parental characteristics for use in research and medical purposes as and when required. Though various protocols have been established to generate organoids from different tissues [16–18], studies on the retention of original tumoral characteristics in the Cervical Cancer OrganoidS (CERCOS) after being cryopreserved is very limited. Therefore, we have also shown that organoids retain the morphological and histopathological features of the parent tissue post cryopreservation.

## Materials and methods

### Patient sample collection

Biopsy samples were collected from patients diagnosed with cervical cancer (squamous / adenocarcinoma) as per the approval of Institute Ethics Committee (IEC no. PGI/IEC/2020/000365). The collection of each sample was supervised by the consulting gynecologist and pathologist.

### Tissue processing

Patient tissue sample was collected in ice-cold PBS and brought to the laboratory within 10 minutes. The sample was washed thoroughly with ice-cold PBS thrice and dissected into small pieces (2-3 mm each) using forceps and a blade. The dissected tissue pieces were then treated with Collagenase I (1mg/ml) for approximately 1-1.5 hours and strained using through 70 μm cell strainer to obtain single cell suspension. The cells were then treated with 5 ml ACK lysis buffer (Gibco) for 5 minutes for RBC lysis. The obtained cell pellet was resuspended in organoid medium (Supplementary table 1) and seeded on Matrigel according to different culture methods followed.

### Matrigel dome method

In matrigel dome method [1], the single cell suspension was resuspended in GFR Matrigel and seeded as drops of 10 μl on pre-warmed 24-well culture plate. The drops were allowed to get solidified for 20 minutes at 37 °C followed by addition of organoid media.

### Matrigel Bilayer Organoid Culture (MBOC) method

The MBOC protocol was followed as described by Maru et al.[2]. Matrigel GFR was pre-solidified on 24-well culture plate and then seeded with single cell suspension. It was incubated overnight to allow the cells to attach. Next day, the floating cells were removed and attached cells were overlaid with another layer of Matrigel. It was allowed to get solidified and then overlaid with media.

### Matrigel Single layer method

In this method, 24-well culture plate was coated with Matrigel GFR. Following the solidification of Matrigel, the single cell suspension was seeded onto coated wells and allowed to attach and proliferate.

### Immunofluorescence staining

The organoids were washed with ice-cold PBS and fixed in 4% PFA for 45 minutes followed by treatment with 0.1% Triton-X for 15 minutes and blocking reagent (1% BSA) for 45 minutes. The organoids were then incubated with phalloidin dye for 1 hour followed by DAPI for 20 minutes. Then the organoids were mounted with anti-fade mountant and visualized using Olympus confocal microscope.

### Scanning Electron Microscopy

Organoids were fixed in 2.5% glutaraldehyde overnight at 4° C followed by PBS wash. The fixed organoids were then dehydrated by washing with series of ethanol gradients (35% to 100%). Then the organoids were air dried and mounted on stub using double stick carbon tape. The organoids were sputter coated with gold and then images were acquired on Jeol – JSM-IT300 Scanning Electron Microscope (SEM).

### Histology and Immunohistochemistry

Histology and immunohistochemistry studies were performed on 10-day old organoids. The organoids were fixed in 4% PFA overnight at 4° C. Then the organoids were embedded in paraffin blocks, sections were cut and hydrated before staining. The sections were then processed for H & E staining and immunohistochemistry by incubation with antibodies against specific marker proteins. Images were captured using Olympus BX53F2 bright field microscope.

### Whole exome sequencing

For whole exome sequencing, DNA from tissue as well as collected organoids was isolated using DNeasy Tissue and Blood Kit respectively (Qiagen) following the manufacturer’s instructions. The isolated DNA was checked for quality on Qubit 4.0 fluorometer (ThermoFisher Scientific USA) before proceeding for library preparation. Library preparation involved enzymatic fragmentation followed by barcode adapter ligation, amplification, magnetic bead-based purification and then target enrichment using exome panel (Twist exome v2.0) and hybridization and amplification mix. Final libraries were quantified by qubit fluorometer and then normalized and loaded on NovaSeq-6000 Flow-cell for PEx150bp and 100x mean depth sequencing. Raw data was demultiplexed and quality checked using FastQC, followed by alignment to reference genome (hg38). The bam files were subjected to duplicate removal using Picard tools and variant calling was performed using HaplotypeCaller (GATK), Mutect-2 and CNVkit. Variant annotation was performed with Annovar followed by variant analysis. The counts for variants and genes common between organoids and tissue and specific to only tissue and organoid respectively were listed using both HaplotypeCaller and Mutect-2. Further, the common genes between organoids and tissue samples were mapped to the cervical cancer genes from CBio Portal to shortlist the important genes related with cervical cancer development. The softwares along with the versions used for different purposes are listed in Supplementary table 2.

### HPV typing

PCR was performed to determine the HPV status of the organoids and the tissue. Infection with any HPV subtype was determined using primers specific for L1 region of the HPV genome. Further, PCR using type-specific primers for two major high-risk subtypes - HPV 16 and HPV 18 was also carried out. GAPDH was used as the house-keeping gene. SiHa and HeLa cell line were used as positive control for HPV 16 and HPV 18 respectively. Details of the primers used can be found in Supplementary table 3.

### Passaging of organoids

To passage organoids, organoid media was removed from the wells and treated with Cell Recovery Solution (Corning) at 4° C for 1 hour. The organoids can be seen floating in the cell recovery solution after one hour. The floating organoids were collected, washed with ice-cold PBS and centrifuged at 570 g for 5 minutes. The obtained organoid pellet was then treated with 1x TrypLE Express enzyme for 5 minutes to break the organoids into small clusters of cells. The obtained cell pellet was then seeded in a ratio of 1:3 in 24-well plates pre-coated with GFR Matrigel.

### Cryopreservation and revival of organoids

The collected organoids were dissociated using TrypLE, followed by resuspending the pellet in the freezing medium (Bambanker) and stored at - 80 °C. The organoids should not be dissociated to single cells. The cryogenic vial with the cryopreserved organoids was incubated for 1-3 minutes at 37 °C. Thereafter, AdDMEM/F12 +++ was added dropwise into the cryogenic vial. The organoids were centrifuged at 370 g for 5 minutes and the pellet was resuspended in organoid medium and seeded on matrigel and incubated at 37 °C and 5 % CO2.

## Results

### Optimization of organoid culture method

Fig. 1 illustrates the utilization of various approaches to enhance the optimization of organoid culture. The Matrigel dome method is widely regarded as the method of choice for organoid culture. However, we did not observe satisfactory organoid formation with this method in any of the patient samples (n = 5) (Fig. 1). Then we tried MBOC protocol which is specifically tailored for organoid culture in the context of gynecological cancers [13–15]. In the case of the MBOC protocol (n=3), organoids begin to form in two of the three samples within a span of just four days; however, they exhibited signs of disintegration shortly thereafter. Then, we modified the MBOC protocol in which only the lower layer of matrigel was used, calling it ‘Matrigel single layer method’. This method yielded the formation of larger-sized organoids (n = 5) within a timeframe of 4 to 7 days, with the potential for proliferation observed up to day 13.

**Fig. 1.**
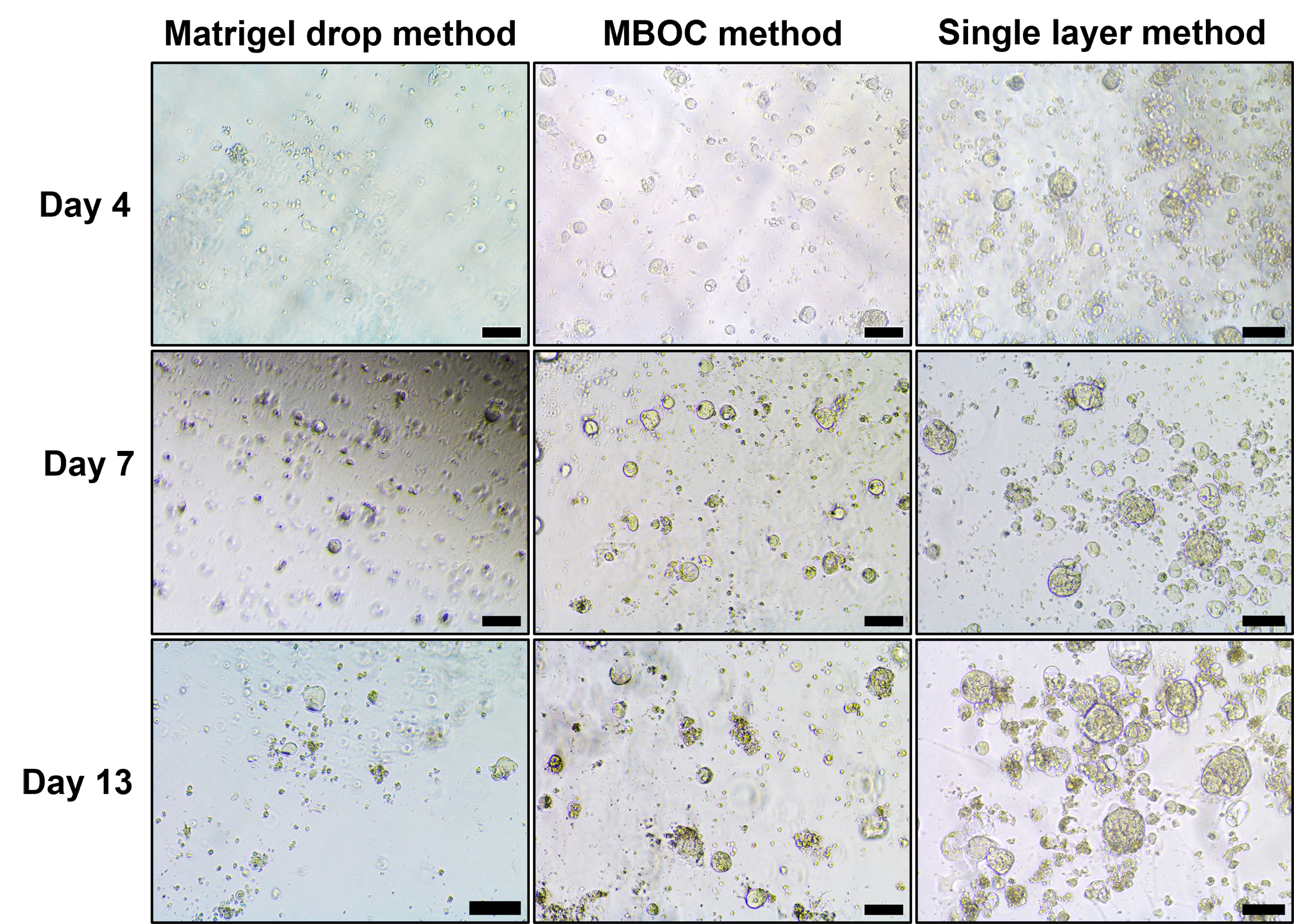
Comparison of different methods for organoid generation. Representative images showing comparison of three different methods (Matrigel dome method, Matrigel Bilayer Organoid Culture method (MBOC) and Matrigel single layer method) employed for organoid generation. Images were taken at different days (day 4, day 7 and day 13); (scale bar, 500 µm).

### Establishment of patient derived organoids (PDOs) using Matrigel single layer method

PDOs were successfully generated from biopsy samples obtained from five patients and were designated as PDO1, PDO2, PDO3, PDO4 and PDO5 (Fig. 2A). The characteristics of the patient samples and the derived organoids are given in Supplementary table 4. PDO1, PDO3, PDO4 and PDO5 were derived from patients diagnosed with cervical squamous cell carcinoma whereas PDO2 was derived from patient diagnosed with cervical adenocarcinoma. The age of the patients ranged from 42 to 50 years old, however one of the patients was 59 years old. We observed variability in the morphology among different PDOs ranging from compact round morphology to grape like morphology. The average size of 7-day old PDOs was observed to be 177 µm. There was also variation in the proliferation potential of the PDOs. PDO2, PDO4 and PDO5 showed good proliferation ability and were successfully characterized and compared with the parent tissue. However, PDO1 and PDO3 could not be characterized due to insufficient amount of sample and limited proliferation potential.

**Fig. 2.**
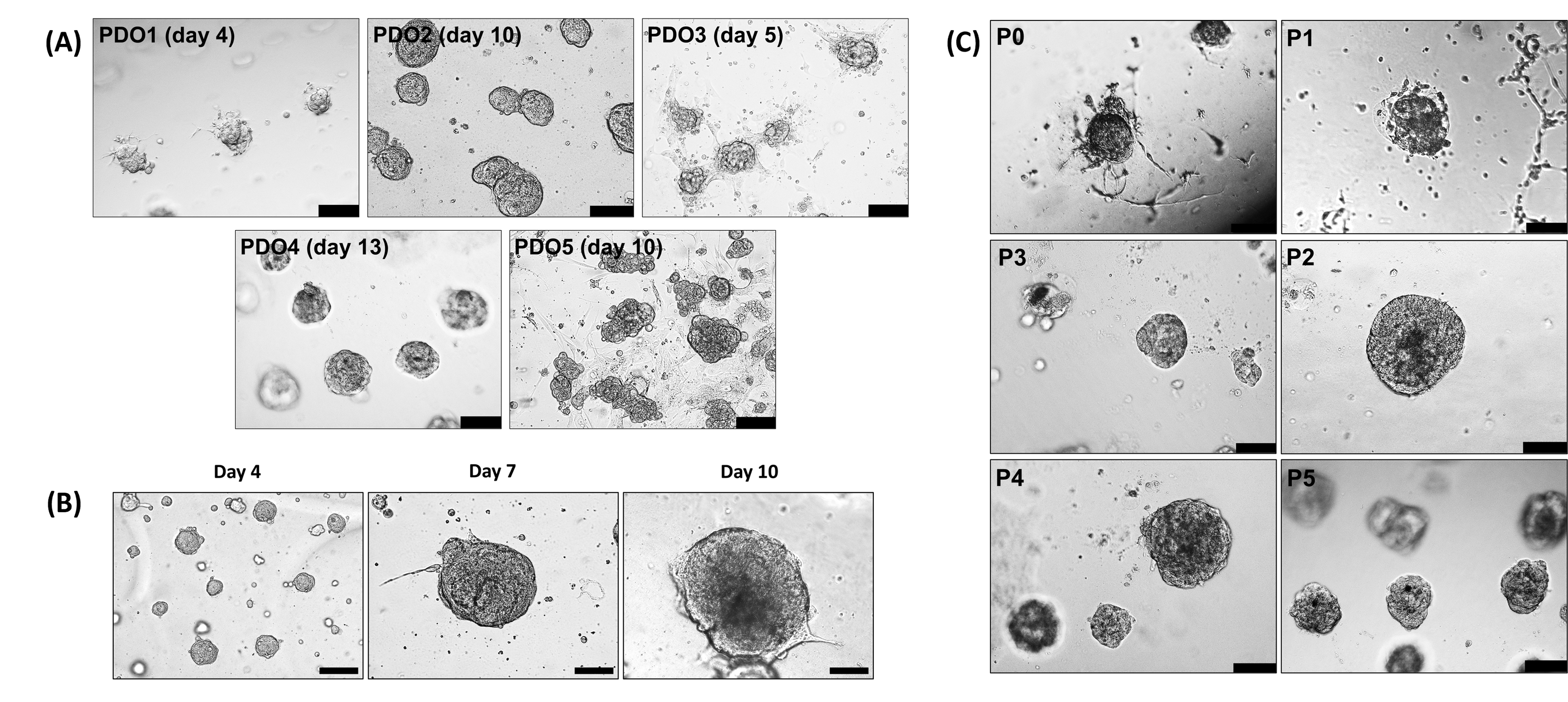
Establishment and expansion of organoids. A. Bright field microscopy showing different morphologies of cervical cancer organoids, from solid to grape like morphology. B. Representative bright field images showing organoid growth (PDO2) at different time points. C. Bright field images of organoids (PDO4) at different passages showing the expansion and proliferation of organoids (Px0 to Px5). (Scale bar, 500 µm)

### Expansion of PDOs

Organoids were passaged after they achieved a size of around 150 μm. It was observed that the ability of organoid expansion varied from sample to sample. We could successfully passage two PDOs for > 2 months (Fig. 2C). PDO4 was grown till fifth passage (P5) and then it was cryopreserved. PDO5 was expanded till passage 4 (P4) before being cryopreserved. The passaged organoids were observed to attain larger size (around 230 μm within 7-10 days). As shown in Fig. 4C, H & E staining of the passaged organoids confirmed the retention of histological features of the original tumor.

### Visualization of actin cytoskeleton

Cytoskeleton plays an important role in 3D structures like organoids and spheroids, and are crucial for maintaining their mechanical integrity. Actin filaments are involved in cell adhesion, cell shape, cell migration and their arrangement facilitate 3D structure formation. To visualize the arrangement of actin filaments in organoids, phalloidin staining was performed. The results of phalloidin staining showed peripheral arrangement of actin filaments (red) with nucleus in the center (blue) (Fig. 3A). This arrangement of actin filaments has been reported to support the organoid formation.

**Fig. 3.**
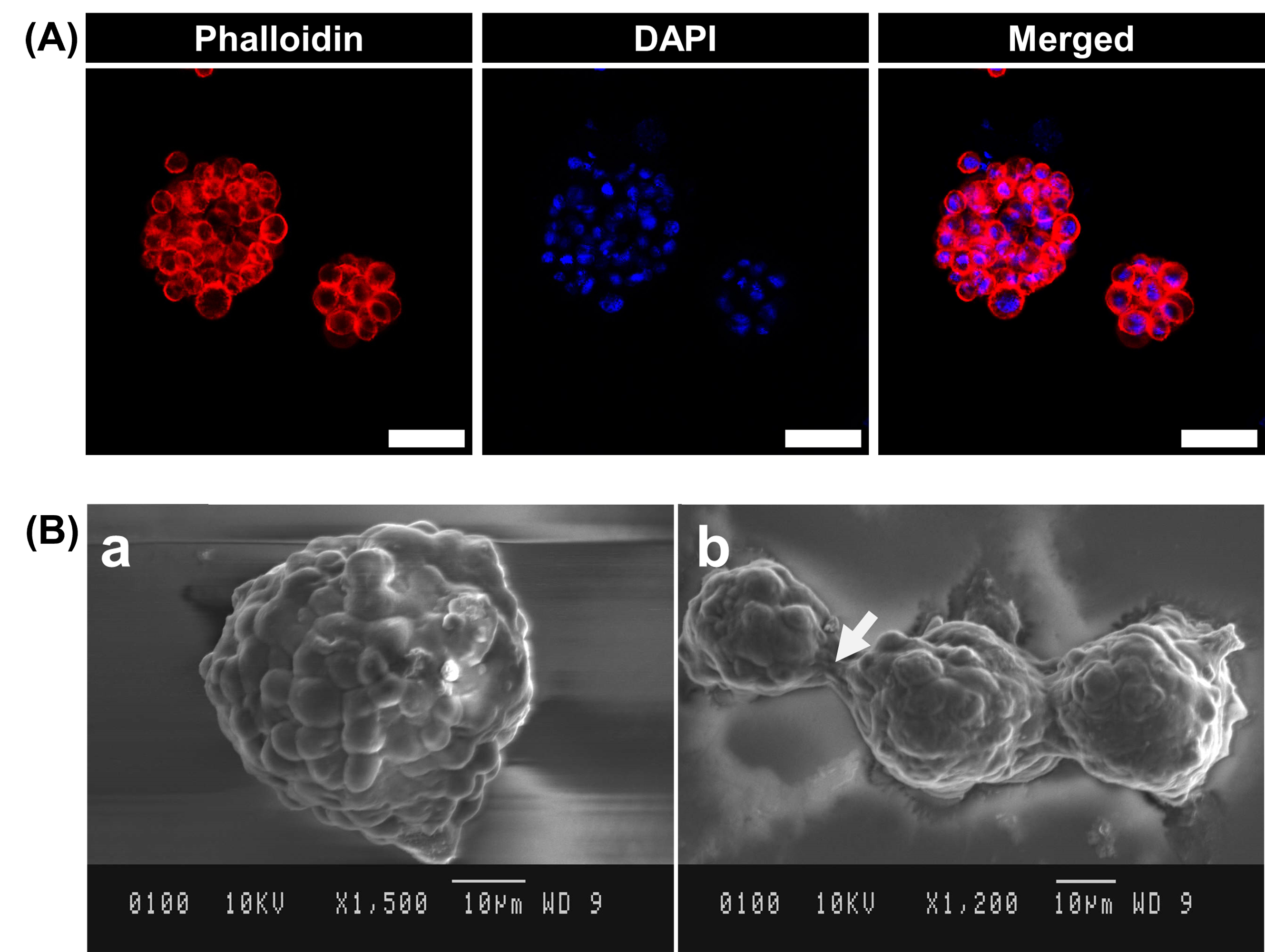
Structural analysis of organoids through phalloidin staining and scanning electron microscopy (SEM). A. Visualization of arrangement of actin filaments using Phalloidin staining (Red – Phalloidin, Blue – DAPI) (scale bar, 50 µm). B. (a) Analysis of ultrastructure of revived PDO4 using scanning electron microscopy (SEM) (x1500). (b) White arrow showing cell-cell connections between PDOs (x1200).

**Fig. 4.**
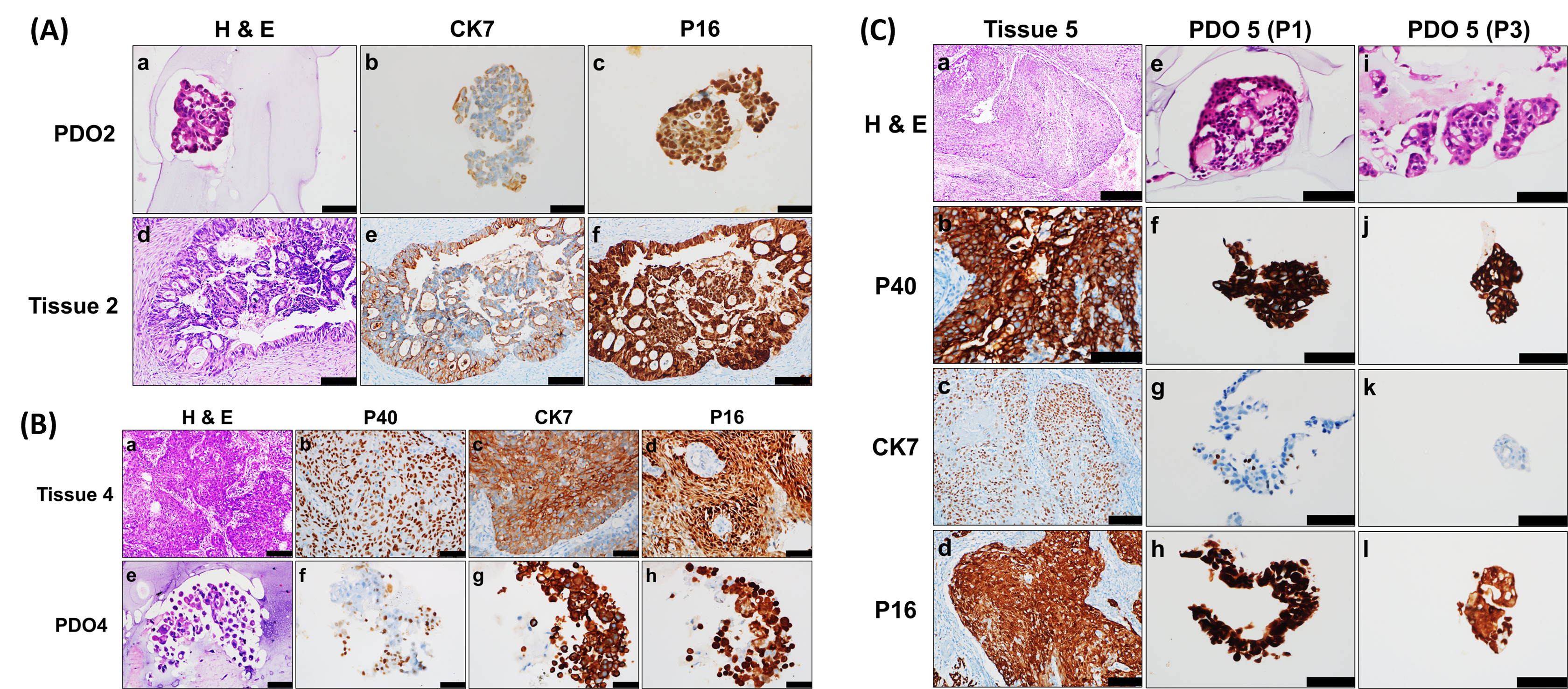
PDOs recapitulate the characteristics of original tumors. H&E stained and IHC-stained images of A. PDO2 - adenocarcinoma; B. PDO4 squamous cell carcinoma and their corresponding tumor tissues. C. H&E stained and IHC-stained images of PDO5 (squamous cell carcinoma) at different passages (Px1 and Px3) and their comparison with the corresponding tumor tissue.

### Analysis of ultrastructure by SEM

SEM imaging of PDOs for ultrastructural analysis showed strong cell-cell interactions forming a compact and smooth profile as illustrated in Fig. 3B. A bulging cell has been observed in some organoids and several short processes, those appearing like microvilli and blebs have also been observed, making the surface irregular [20–21].

### Histology and immunohistochemical analysis

Histological and immunohistochemical analyses were conducted to compare PDOs with corresponding patient tissues. PDO2 showed a glandular cell arrangement consistent with adenocarcinoma, while PDO4 and PDO5 exhibited similar cell arrangements to the squamous cell carcinoma subtype of the original tissues. p16 expression, a surrogate marker for HPV infection, and CK7, an epithelial biomarker, were assessed in all PDOs and tissues. p40, specific for squamous cell carcinoma, was only evaluated in squamous carcinoma samples. IHC analysis revealed strong concordance in p16 and CK7 expression between PDOs and original tumors, confirming tissue recapitulation. p40 expression was similarly observed in PDO4, PDO5, and their corresponding tumors.

### Whole Exome Sequencing data from tissues and corresponding organoids

Whole exome sequencing of 3 tissues (Tissue 2, Tissue 4 and Tissue 5) and their corresponding PDOs (PDO2, PDO4 and PDO5) was done at 100x depth and variant calling was done using Mutect2.

Mutect2 identified a total of 340404, 391516 and 423218 variants in Tissue 2, Tissue 4 and Tissue 5 respectively. A total of 549290, 429588 and 389839 variants were identified in PDO2, PDO4 and PDO5 respectively. An average of 385016 identified in the three tissues and 456239 variants identified in the three PDOs. Out of these, 95224, 83947 and 97067 common variants found between tissue 2 and PDO2, tissue 4 and PDO4, and tissue 5 and PDO5 respectively (Supplementary Fig. 1).

### Common variants related to cervical cancer development

The identified common variants between patient tissue and corresponding PDO were then uploaded to CBioPortal. All the variants and associated genes reported in cervical cancer related studies and present in CBioportal even at a very low allele frequency were taken (Supplementary table 5). Further filtering was performed and variants with allele frequency ≥ 0.01 in population databases (ExAc), low coverage depth (<5 total reads), less than 2 variant allele reads and benign computational prediction by computational tools (SIFT and Mutation Taster) were excluded. The remaining variants were categorized as per ACMG and AMP criteria. Cervical cancer tissues were found to harbor a higher number of variants compared to their paired organoids (416 vs. 150; p = 0.0115), likely due to the selective growth of less aberrant cell subpopulations in culture and/or the presence of low-frequency subclones in the original tumors (Supplementary Fig. 2). The filtered variants were then mapped with the cervical cancer related genes taken from CBioportal and Cosmic database. The results showed that PDO2 and PDO5 shared 50% similarity with the paired tissue whereas PDO4 shared 31% similarity with the paired tissue (Fig. 5A). Among the top common mutated genes were *LRP1B, KMT2A, MET, TTN, PIK3CA* and *NF1*. Mutations in some cancer related genes such as *MTOR, ALK, KMT2C* and *KMT2D* were found to be lost in the derived organoids. The genes which were found to be mutated in more than one sample were SMO, TTN and VEGFA. We further analyzed base variants in cervical cancer tumor and organoids. The point variant type of parent tissue was well conserved in the derived organoids with C > T being the dominant base variant type (Fig. 5B).

**Fig. 5.**
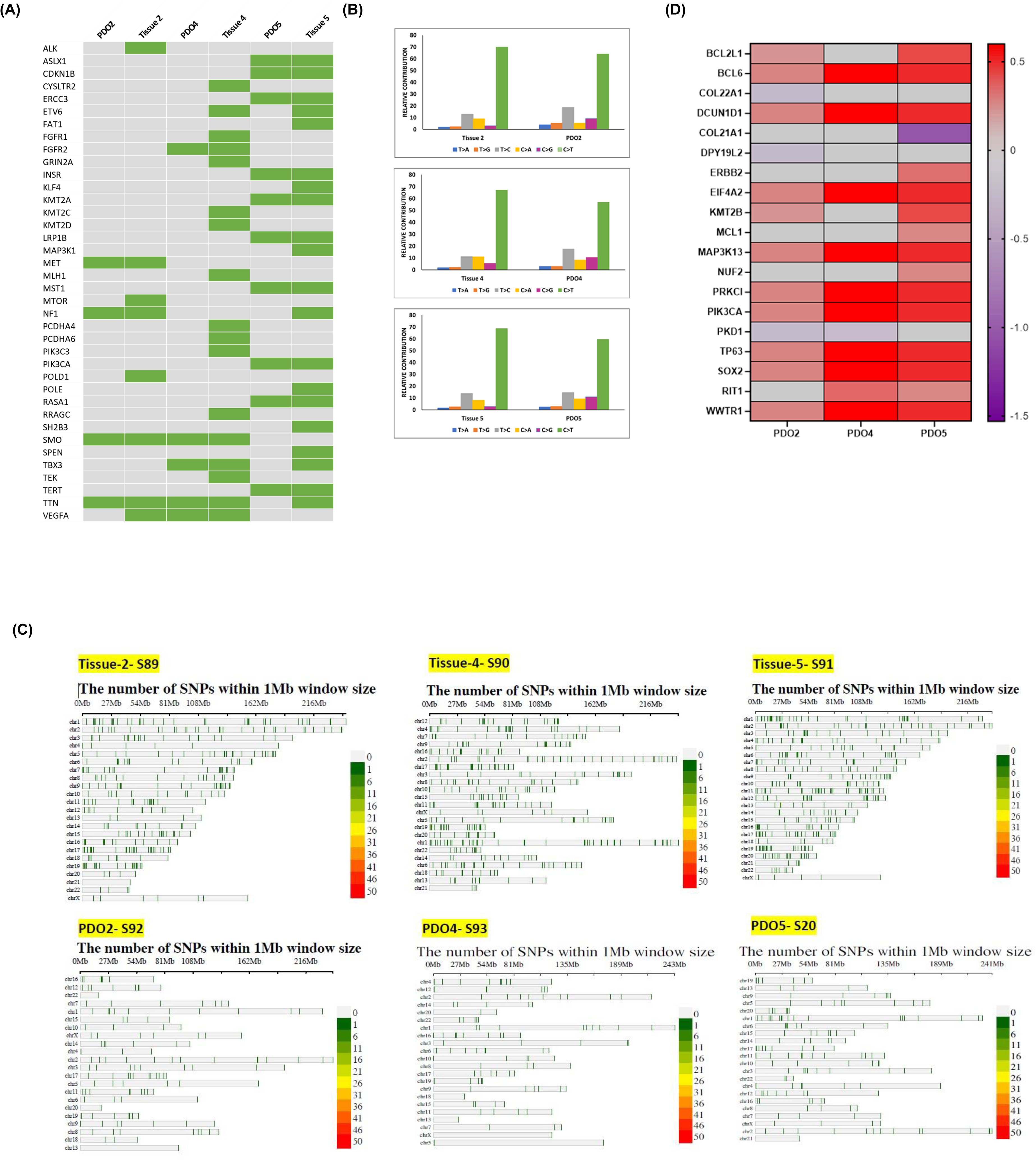
Comparison of mutational landscape of organoids and the corresponding tissues through WES. A. Mutation matrix comparing the somatic mutated cancer genes in cervical cancer organoids with those in parent tumor tissues. B. Bar plots showing that frequency of six types of point mutation types in the parent tissue and the derived organoids with C > T being the dominant SNV type. C. Heat map showing the CNVs (log fold change) in the cervical cancer related genes in organoids compared to the corresponding tumor tissue. (D) SNP density plots showing reduction in SNP density in PDOs compared to corresponding tissues

We have also plotted single nucleotide polymorphism (SNP) density plots of tissues and PDOs post applying filtering criteria. It was observed that there was a reduction in the SNP density in the PDOs compared to the corresponding tissues (Fig. 5C).

### Copy Number Variation (CNV) Calling

CNVkit tool was used for CNV calling in the tissues and the PDOs. The tissue samples were taken as the reference to calculate the log fold change in CNVs in the corresponding PDOs (Supplementary table 6).

Top 20 genes showing CNV gain in cervical cancer according to Cosmic database were found to be amplified in the three PDOs and all the genes were found to be localized on chr 3. On comparing copy number alterations present in CBioportal with frequency >10%, 25 genes were found to exhibit fold change in CNV in PDO2, 31 genes in PDO4 and 36 genes in PDO5. Among them, *ATR* (3q23), *SOX2* (3q26.33), *BCL6* (3q27.3), *EIF4A2* (3q27.3), *PRKCI* (3q26.2), *PIK3CA* (3q26.32), *TP63* (3q28), *DCUN1D1* (3q26.33), *FGF12* (3q28-q29) and *MAP3K13* (3q27.2) were among the most common amplified genes (Fig. 5D).

### HPV typing

HPV typing using PCR showed that organoids could recapitulate the HPV status of the organoids. PDO2 and the corresponding tissue showed positivity for L1 region indicative of HPV infection and HPV 18. However, PDO2 was negative for HPV 16 whereas the corresponding tissue was positive for HPV 16. PDO4 and the corresponding tissue were positive for L1 and HPV 16. PDO5 and the corresponding tissue showed positivity for L1, HPV 16 and HPV 18 (Supplementary Fig. 3).

### Cryopreservation and revival of organoids

PDO4 was cryopreserved and revived after one month. Initially, there was slow proliferation of the revived organoids. It took around 11 days to proliferate and then it was passaged. Following the passage, there was an increase in proliferation rate as the organoid was formed within 7 days. Morphological analysis of the revived organoids using SEM showed compact and round morphology, indicating preservation of cell-cell connections (Fig. 6C). Histological analysis showed preservation of histological features, indicative of its squamous cell carcinoma type (Fig. 4(B) and 6(B)). Further, IHC analysis of revived organoids showed similar expression of diagnostic markers p63 and CK7 as in original tissue, indicating preservation of histological subtype even post cryopreservation (Fig. 4B and 6B).

**Fig. 6.**
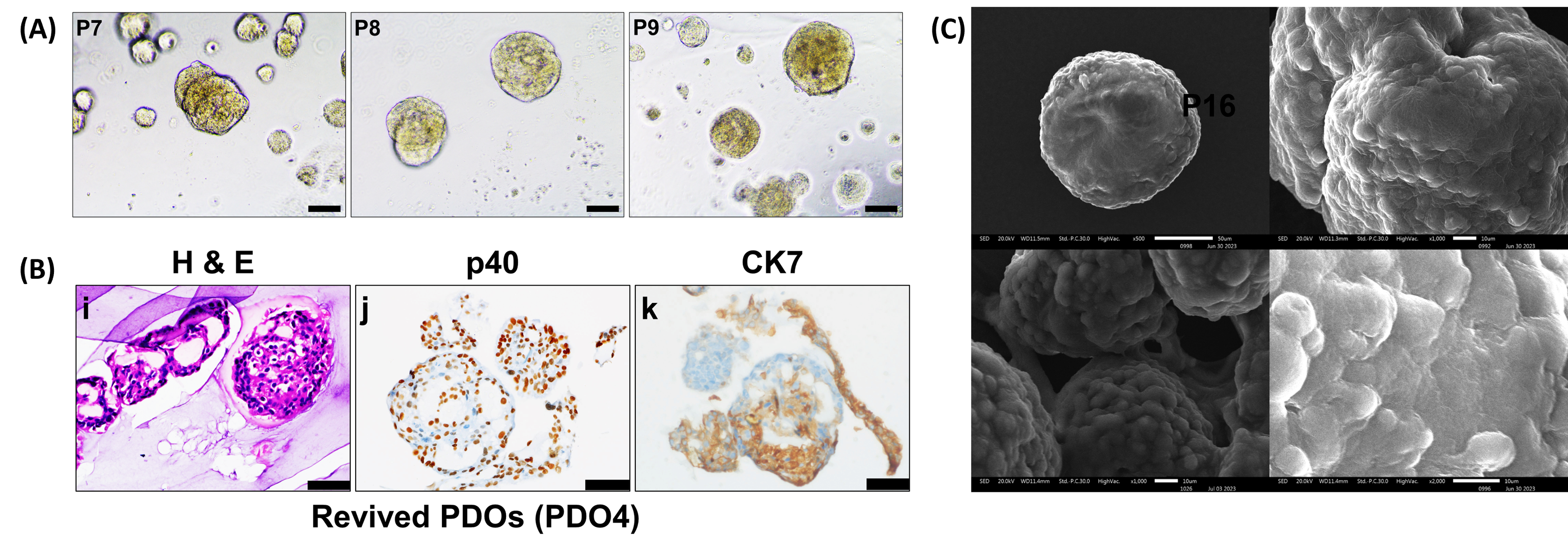
Characterization of PDOs post cryopreservation and revival. A. Bright field images of PDO4 at different passages (Px6, Px7 and Px8) post cryopreservation and revival (Scale bar, 500 µm). B. Histological and IHC analysis of revived PDOs showing morphology and expression of cell surface markers (P40 and CK7) post cryopreservation (Scale bar, 100 µm). C. Analysis of ultrastructure of revived PDO4 using scanning electron microscopy (SEM) (x500, x1000 and x2000).

## Discussion

PDO models have been reported to successfully recapitulate the intra- and inter-tumor heterogeneity. These models serve as a bridge between conventional in vitro models and in vivo models and have immense potential for clinical applications, particularly in the field of cancer PDO models mimic the tumor tissue phenotype more closely compared to conventional 2D cultures. Studies now demonstrate a matched drug response between the organoids and the corresponding patients making then a suitable platform to predict the chemotherapy or targeted therapy response in many of these cancer [16,18,22–24].

In the present study, we optimized the organoid cultures and observed that our simple modification of the Matrigel monolayer method was useful for efficiently establishing organoids from three cervical cancer types that contain different subtypes. Matrigel dome method has been widely adopted for organoid generation of various cancers including breast cancer, lung cancer, pancreatic cancer, ovarian cancer and rectal cancer [13–17,25]. However, in the present study, Matrigel dome method didn’t produce satisfactory organoids from cervical cancer tissue even after 14-20 days, which is the usual period for organoid generation reported for this method. This is probably due to the accumulation of detrimental factors released from dead cells trapped within the dome as reported previously [13]. Modified Matrigel Bilayer Organoid Culture (MBOC) protocol has been reported to capture highly proliferative stem cell like cell populations without conducting any cell-sorting in gynecological tumors [26], therefore this method was employed for organoid formation. Though organoids were formed within 4-7 days using MBOC protocol, but these disintegrated soon. We concluded that the presence of upper layer of Matrigel might be the cause that limited the effective organoid proliferation. Consequently, only the lower layer of Matrigel was utilized, and this variation in the protocol was termed the ’Matrigel Single Layer Method’. This method generated comparatively large sized organoids (∼150 μm) in a smaller number of days (generally 7-10) which were robust and could be passaged. Therefore, this modified method of cervical organoid generation was adopted for further characterization.

Our results show that PDOs could be successfully recovered from the matrigel after culturing with matrigel single layer method. In our study, matrigel were depolymerized with cell recovery solution and the organoids were collected without any vigorous pipetting or scrapping that often compromises the viability of PDOs. This method has an advantage over Matrigel dome method where scrapping and pipetting is sometimes required to release the organoids from the dome [27–28].

Histological characterization and profiling of the relevant biomarkers (p16, p40 and CK7) demonstrated the preservation of key features of the corresponding tumor biopsy. Further PDOs retained the characteristics of HPV genome of the parent tissue as both tissues and PDOs showed similar expression of HPV 16 and HPV 18. This indicates that the organoid models may serve as a suitable platform to further understand the pathophysiology of HPV in the host.

An important aspect of the study was that PDOs could be cryopreserved and revived efficiently. The cryopreservation was done using a simple and robust method involving resuspension of cell pellet in the freezing medium [17]. It was observed that the cryopreserved PDO4 retained the morphological and histological characteristics of the parent tumor. This demonstrated that PDOs can be cryopreserved in an organoid biobank and can be used for molecular profiling, drug testing, and biomarker discovery for personalized medicine. .

One of the important features of organoids is that they maintain the mutational landscape of the parent tumor. PDO2 and PDO4 showed a few common mutations with their parent tissues, while PDO5 had more shared somatic variants. Some mutations were lost in PDOs, likely due to the procedures of tumor tissue sampling and organoid culture development. Additionally, inherent tumor characteristics such as purity, stage, and regional heterogeneity—may have influenced organoid outgrowth efficiency. These factors could lead to the loss of specific tumor sub-clones, ultimately resulting in tumor–organoid pairs with low concordance . The most frequent mutation identified was in the titin (TTN) gene, present in all three patient tissues and two organoids (PDO2 and PDO4), but absent in PDO5. TTN plays a crucial role in cardiac and skeletal muscles and is associated with increased mutation load, improved response to immune checkpoint therapy, and longer survival in various solid tumors, including cervical cancer [30]. Another commonly mutated cervical cancer gene in squamous cell carcinoma (28 %) is PIK3CA with E545K mutation being a major hotspot mutation [31]. This mutation causes abnormal cell proliferation and reduced apoptosis, which has been linked to cervical cancer development. The other mutated gene was Smoothened (SMO), a conserved signal transducer of the Sonic Hedgehog pathway. Various studies have reported the role of Hedgehog signaling pathway in mediating chemo-radio resistance and regulating stem cell characteristics during EMT transition in cervical cancer [32–35]. High CNV fold change in organoids compared to the original tumor can be attributed to the high proliferative potential of stem cells forming the organoids and the manipulation of *in vitro* culture conditions by the addition of external growth factors [36–38].

In conclusion, the present study illustrates the successful establishment and characterization of PDOs through a modified and time-efficient culture system. The developed PDOs were observed to preserve the tumor heterogeneity while retaining the histopathological and genetic features of the original tumor. The histopathological features were observed to be retained even post cryopreservation. This suggests that the cervical cancer PDOs may be employed as a robust platform for *in vitro* drug screening a robust platform for personalized medicine validation and treatment plan optimization.

## Acknowledgments

SB acknowledges the Indian Council of Medical Research (ICMR) for providing extra mural funding for the study (grant ID 2020-4349). SS acknowledges Council of Scientific and Industrial Research (CSIR) for providing research fellowship. The authors are grateful to CSIC facility, PGIMER for carrying out Scanning Electron Microscopy.

## Author contributions

Conceptualization: SS, SB; Data curation: SS, RB, RS, PB, SB; Formal analysis: RB, RS, PB, SB; Funding acquisition: SB; Investigation: SS, RB, RS, SB; Methodology: SS, PB, SB; Project administration: SB; Resources: SB; Software: PB; Supervision: RB, RS, PB, SB; Validation: SS, RB, RS, PB, SB; Visualization: PB, SB; Roles/Writing - original draft: SS; and Writing - review & editing: RB, RS, PB, SB.

## Data Availability Statement

Data will be available on request.

## Ethics Statement

Ethical approval was obtained from the institute for carrying out the study (IEC no. PGI/IEC/2020/000365). The samples were collected after obtaining written consent from all the patients.

## Abbreviations

CK7: Cytokeratin 7
CNV: Copy Number variation
GFR Matrigel: Growth Factor Reduced Matrigel
HPV: Human Papillomavirus
MBOC: Matrigel Bilayer organoid culture
PDO: Patient-derived organoids
SEM: Scanning Electron Microscope
SNP: Single Nucleotide Polymorphism

