## Supplementary figures and images for "Developing *in vitro* patient derived CERvical Cancer OrganoidS (CERCOS) as a potential preclinical tool in cervical cancer research"

### Suplementary Fig. 1

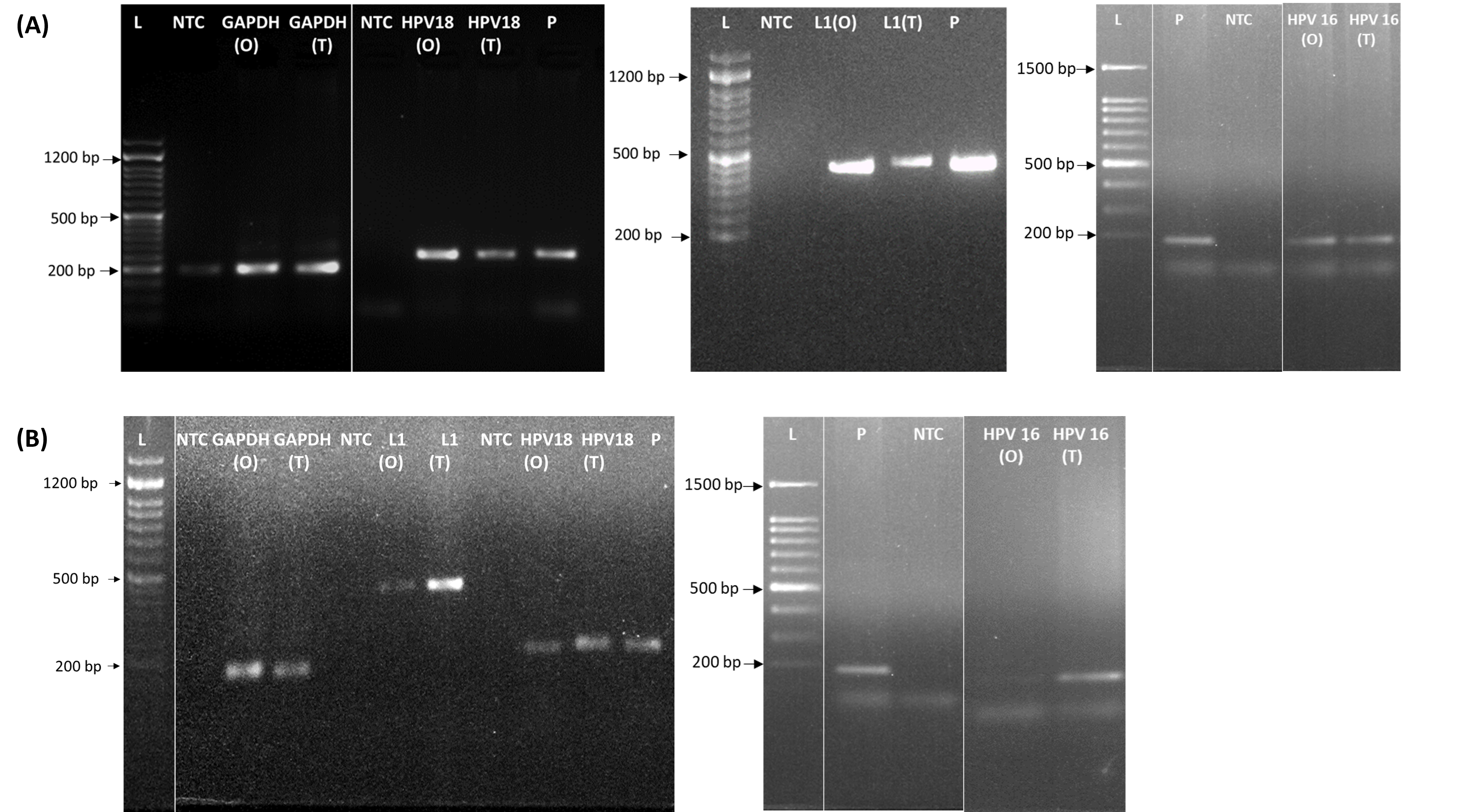

### Suplementary Fig. 2

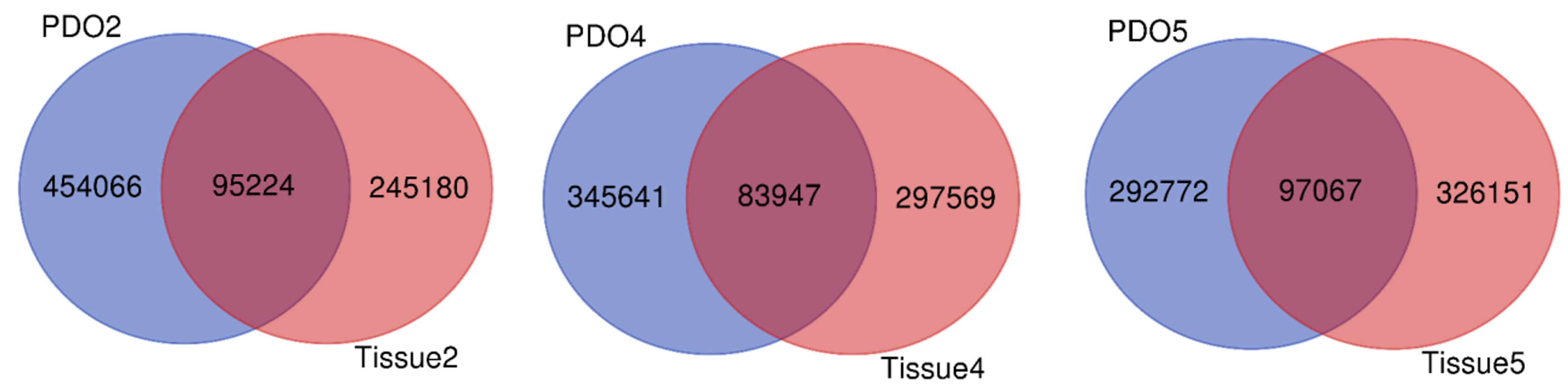

### Suplementary Fig. 3

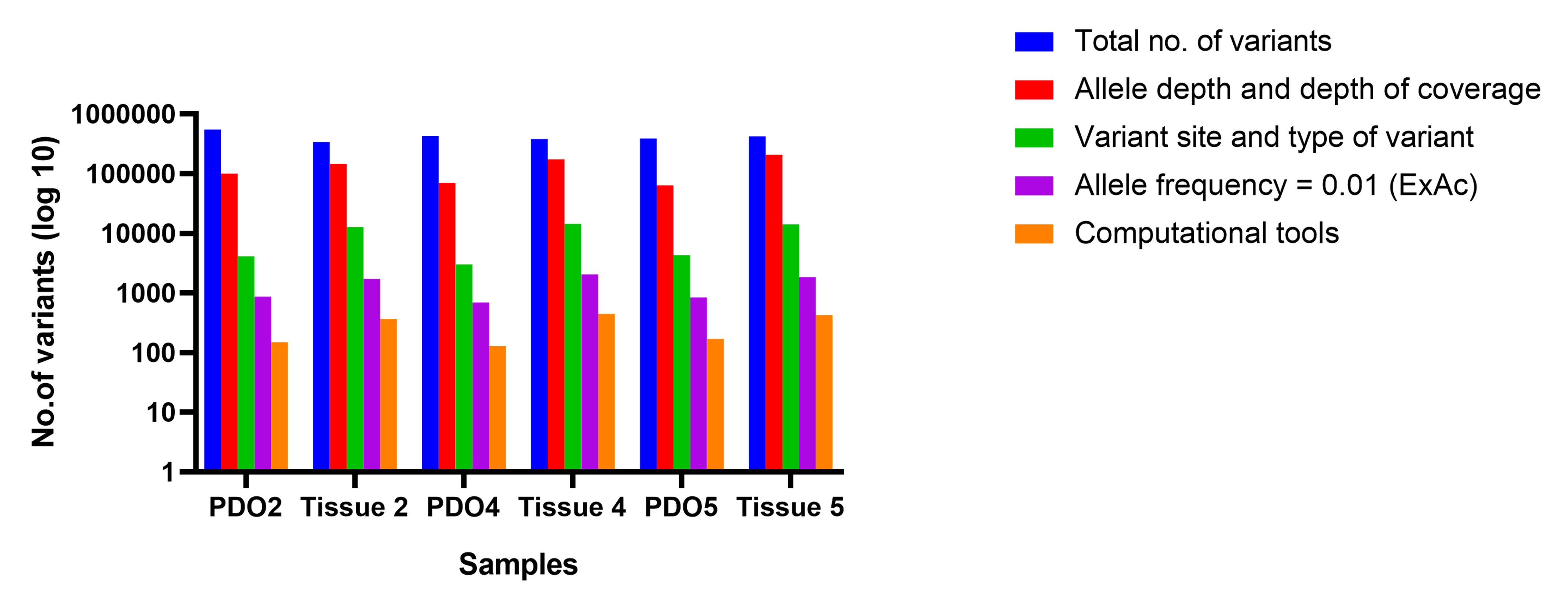

### Suplementary table 6

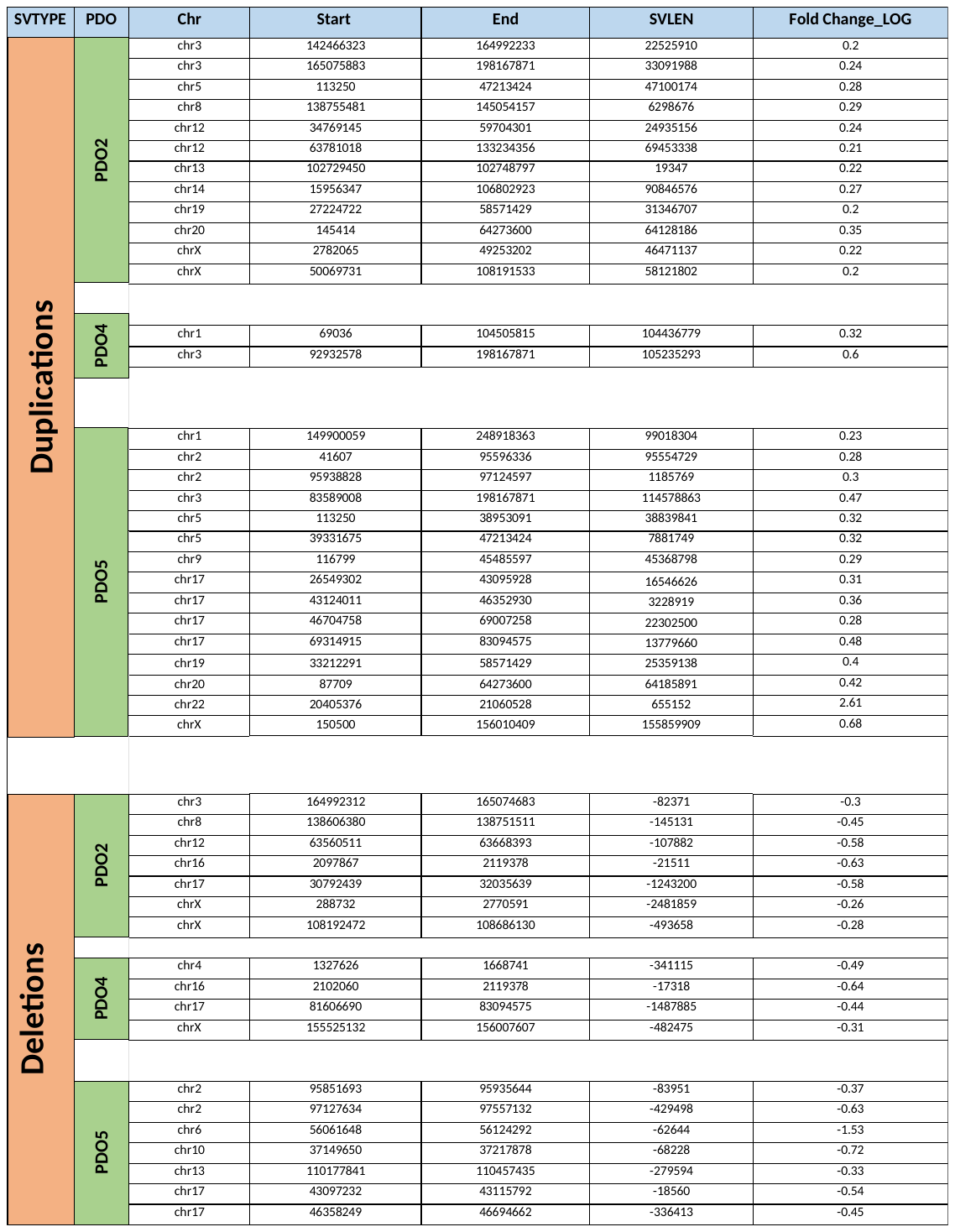
