## Supplementary material for "Developing *in vitro* patient derived CERvical Cancer OrganoidS (CERCOS) as a potential preclinical tool in cervical cancer research": Suplementary table 1

|  | **Component** | **Conc. used** |
| --- | --- | --- |
| 1 | AdDMEM/F12 (Gibco) |  |
| 2 | Glutamax (Gibco) | 1x |
| 3 | HEPES (Sigma) | 1 mM |
| 4 | Penicillin-streptomycin-amphotericin solution | 1x |
| 5 | B27 (Gibco) | 1x |
| 6 | Rock inhibitor (Y27632) (Abmole) | 10 µM |
| 7 | Jagged-1 (Anaspec) | 1 µM |
| 8 | bFGF (Gibco) | 20 ng/ml |
| 9 | Recombinant Human Noggin (Peproetch) | 100 ng/ml |
| 10 | Recombinant Human R-spodnin 1 (R & D systems) | 250 ng/ml |
| 11 | EGF (Gibco) | 50 ng/ml |
| 12 | Nicotinamide (Sigma) | 2.5 mM |
| 13 | N-acetyl-L-cysteine (Sigma) | 1.25 mM |
| 14 | A83-01 (Tocris) | 500 nM |
| 15 | p38 inhibitor SB2020190 (Sigma) | 1 µM |
