## Supplementary material for "Developing *in vitro* patient derived CERvical Cancer OrganoidS (CERCOS) as a potential preclinical tool in cervical cancer research": Suplementary table 2

| **Process Names** | **Software** | **Versions** |
| --- | --- | --- |
| BCFTOOLS_SORT | bcftools | 1.17 |
| BCFTOOLS_STATS | bcftools | 1.17 |
| BWAMEM1_MEM | bwa | 0.7.17-r1188 |
| samtools | | 1.16.1 |
| CRAM_TO_BAM | samtools | 1.17 |
| CREATE_INTERVALS_BED | gawk | 5.1.0 |
| CUSTOM_DUMPSOFTWAREVERSIONS | python | 3.11.0 |
| yaml | | 6 |
| ENSEMBLVEP_VEP | ensemblvep | 108.2 |
| FASTP | fastp | 0.23.4 |
| FASTQC | fastqc | 0.11.9 |
| FREEBAYES | freebayes | 1.3.6 |
| GATK4_MARKDUPLICATES | gatk4 | 4.4.0.0 |
| samtools | | 1.17 |
| INDEX_MERGE_BAM | samtools | 1.17 |
| MERGE_BAM | samtools | 1.17 |
| MERGE_FREEBAYES | gatk4 | 4.4.0.0 |
| MOSDEPTH | mosdepth | 0.3.3 |
| SAMTOOLS_STATS | samtools | 1.17 |
| TABIX_BGZIPTABIX_INTERVAL_COMBINED | tabix | 1.12 |
| TABIX_BGZIPTABIX_INTERVAL_SPLIT | tabix | 1.12 |
| TABIX_TABIX | tabix | 1.12 |
| VCFTOOLS_TSTV_COUNT | vcftools | 0.1.16 |
| Workflow | Nextflow | 23.04.2 |
| nf-core/sarek | | 3.2.3 |

**Supplementary table 2**: Details of the software along with the versions that has been used for whole exome sequencing.
