## Supplementary material for "Developing *in vitro* patient derived CERvical Cancer OrganoidS (CERCOS) as a potential preclinical tool in cervical cancer research": Suplementary table 3

| **Primer region** | **Primer sequence** | **Annealing temperature** |
| --- | --- | --- |
| **HPV 16** | F - TGAGCAATTAAATGACAGCTCAGAG | 55 °C |
|  | R - TGAGAACAGATGGGGCACACAAT |  |
| **HPV 18** | F - GACCTTCTATGTCACGAGCAATT | 55 °C |
|  | R - GACCTTCTATGTCACGAGCAATT |  |
| **L1** | F - CGTCCMARRGGAWACTGATC | 55 °C |
|  | R - GCMCAGGGWCATAAYAATGG |  |
| **GAPDH** | F - GGAGCGAGATCCCTCCAAAAT | 55 °C |
|  | R - GGCTGTTGTCATACTTCTCATGG |  |
