## Supplementary material for "Developing *in vitro* patient derived CERvical Cancer OrganoidS (CERCOS) as a potential preclinical tool in cervical cancer research": Suplementary table 4

| **Patient Id** | **Age** | **Cervical cancer subtype** | **Sampling method** | **Samples** | **Establishment and characterization** | **Histopathological features** | | | **Cryopreservation** |
| --- | --- | --- | --- | --- | --- | --- | --- | --- | --- |
|  | | | | | | **P16** | **CK7** | **P40** |  |
| P1 | 43 | Squamous Cell Carcinoma | Punch Biopsy | Tissue 1 | Established, not characterized | NA | NA | NA | No |
|  |  |  |  | PDO1 |  | NA | NA | NA |  |
| P2 | 50 | Adenocarcinoma | Punch Biopsy | Tissue 2 | Established, well characterized | Positive | Positive | NA | No |
|  |  |  |  | PDO2 |  | Positive | Positive | NA |  |
| P3 | 60 | Squamous Cell Carcinoma | Punch Biopsy | Tissue 3 | Established, not characterized | NA | NA | NA | No |
|  |  |  |  | PDO3 |  | NA | NA | NA |  |
| P4 | 49 | Squamous Cell Carcinoma | Punch Biopsy | Tissue 4 | Established, well characterized | Positive | Positive | Positive | Yes |
|  |  |  |  | PDO4 |  | Positive | Positive | Positive |  |
| P5 | 42 | Squamous Cell Carcinoma | Punch Biopsy | Tissue 5 | Established, well characterized | Positive | Positive | Positive | Yes |
|  |  |  |  | PDO5 |  | Positive | Slight Positive | Positive |  |

**Table 1:** Characteristics of patient samples and the derived organoids.
